# Horizontally transferred gene clusters in *E. coli* match size expectations from uber-operons

**DOI:** 10.1101/041418

**Authors:** Tin Yau Pang, Martin J. Lercher

## Abstract

Adaptation of bacteria occurs predominantly via horizontal gene transfer. While it is widely recognized that horizontal gene acquisitions frequently encompass multiple genes, it is currently unclear what the size distribution of successfully transferred DNA segments looks like and what evolutionary forces shape this distribution. Here, we identified 7,538 gene pairs that were consistently co-gained on the same branches across a phylogeny of 53 *E. coli* strains. These pairs are significantly enriched in genes that share the same GO annotation. We estimated the genomic distances of these co-gained pairs at the time they were transferred to their host genomes, which shows a sharp upper bound at 30kb. This upper bound is significantly lower than the size limit on gene co-transfers imposed by the carrying capacity of the transfer agents. The observed distance distribution also appears inconsistent with a model based on the co-transfer of genes within operons; instead, we found that the distance distribution of co-transferred genes closely matches the distribution expected from the transfer of uber-operons, i.e., genomic clusters of co-functioning genes beyond operons.

## INTRODUCTION

Bacterial adaptation to changes in the environment often occurs through horizontal gene transfer (HGT) (Pál et al. 2005; Soucy et al. 2015), i.e., the uptake of genes from genomes of other strains or even other species. Bacteria can exchange DNA through diverse mechanisms including transformation, transduction, conjugation, gene transfer agents, and nanotubes (Davison 1999; Dubey and Ben-Yehuda 2011). If the incoming DNA sequence is highly similar to sequences of the recipient bacterium, then it can be integrated via homologous recombination (Dixit et al. 2015). Otherwise, the foreign DNA segments may be added to the genome through non-homologous recombination after entering the host, resulting in HGT. If transferred genes confer phenotypic changes that provide fitness advantages, then they are likely to become fixed in the bacterial population.

Bacterial genomes are highly dynamic (Soucy et al. 2015). In addition to gene acquisitions via HGT, gene losses via mutational deletions are also frequent among bacteria, a process accelerated by a mutational bias towards gene deletions (Batut et al. 2014); genes no longer needed in the current environment(s) will thus eventually get lost from bacterial genomes. The local pan-genome, the union of all genes in the environment, can be viewed as a toolbox of genes, and HGT often allows bacteria to acquire the genes needed for adaptation from this toolbox (Pál et al. 2005; Maslov et al. 2009; Pang and Maslov 2011). Many phenotypes require the cooperation of two or more genes; accordingly, the joint presence or absence of two genes across many genomes can be used to identify functional associations between them, a method termed phylogenetic profiling (Pellegrini et al. 1999).

Functionally related genes are often co-expressed from the same operon. While HGT may not be the driving force behind operon formation (Price et al. 2005), the operon structure of bacterial genomes facilitates HGT, as it allows the co-transfer of a group of co-functional genes by concentrating them on a relatively small continuous stretch of DNA (Lawrence and Roth 1996). There is also anecdotal evidence for the existence of uber-operons, clusters of functionally related genes in prokaryotic genomes that extend and persist beyond co-transcribed operons (Lathe et al. 2000), and we hypothesized that such larger units may also contribute to the co-transfer of interacting genes.

Previous work has established functional and genomic clustering of co-transferred gene pairs. A systematic analysis of horizontally transferred metabolic genes in proteobacteria confirmed that co-transferred gene pairs are indeed five times more likely to function in the same pathway compared to separately transferred genes (Dilthey and Lercher 2015). The same study also found that co-transferred gene pairs are more than twice as likely as random pairs to be genomic neighbours (defined as genes separated by at most two intervening genes) (Dilthey and Lercher 2015).

To test if operons or uber-operons are the basic units of HGT, we reconstructed the phylogenetic tree of 53 *E. coli* and *Shigella* strains, and identified gene pairs that were consistently co-gained (or co-lost). While we found that *E. coli* operons are too small to explain the observed distance distribution of co-transferred genes, expectations from uber-operons closely match the empirical distribution. These findings show that HGT in *E. coli* is not constrained by the carrying capacities of transfer agents, but by the size distribution of functional gene clusters beyond operons.

## RESULTS

We identified orthologous gene families across 53 *E. coli* and *Shigella* strains (along with 17 strains of other species that served as the outgroup; Supplemental Table S1). *Shigella* strains are generally considered to belong to the species *E. coli* (Chaudhuri and Henderson 2012); thus, we will subsume all 53 strains under the species name *E. coli* in the remainder of this paper. We reconstructed a maximum-likelihood phylogeny based on the concatenated alignment of 1,334 1-to-1 orthologs universally present in all 70 genomes. The resulting rooted *E. coli* phylogeny, which represents vertical inheritance among the 53 strains, is well supported: each internal branch was retrieved in at least 60% of bootstrap samples (see Supplemental Figure S1 for the *E. coli* tree, and Supplemental Figure S2 for the tree including outgroup strains). Based on the assumption that gains and losses are rare events (Pál et al. 2005), we used a maximum-parsimony algorithm on gene presence/absence (Kunin and Ouzounis 2003) to identify gene losses and gains (HGT) along the phylogenetic branches (Methods).

### Statistical association of gene pairs across transfer events

For each pair of orthologous gene families (each “gene pair”) in our dataset, we calculated scores for associations between gains and losses of the two genes across the 104 branches of the *E. coli* tree (Methods). There are three types of pairwise associations: (i) repeated co-gains of two genes via HGT; (ii) repeated co-losses of two genes; and (iii) repeated associations between the gain of one gene and the loss of the other. Co-gained and co-lost pairs may indicate a functional co-operation of the genes. Conversely, the consistent association of the gain of one gene with the loss of another gene (non-homologous replacement) may indicate functional redundancy of the two genes. While many co-gained pairs likely occur through the simultaneous acquisition of both genes on one DNA segment, some co-gained pairs could also stem from distinct HGT events.

We compared the distribution of association scores (aggregated over co-gained, gained-lost, and co-lost pairs) for all gene pairs in the empirical data with that of a null model based on randomizations (Figure 1). The score distributions for the empirical data and the random null model are significantly different, indicating that some gene pairs show much more co-gains, co-losses, or non-homologous replacements than expected by chance.

**Figure 1.**
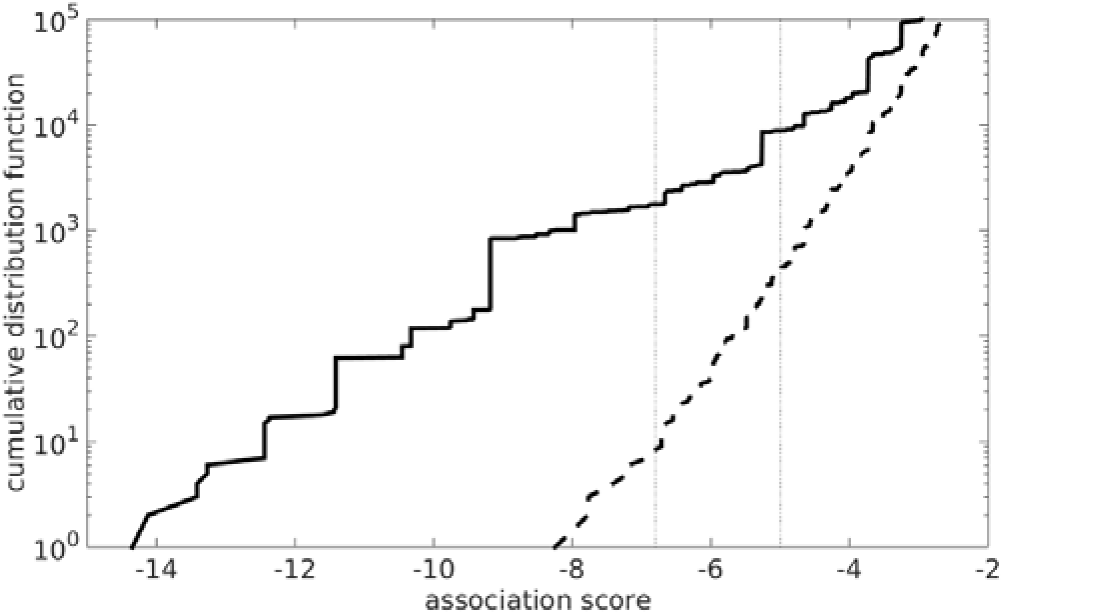
Distribution of the score for pairwise gene associations in the empirical data (solid line) and the null model (dashed line). Each line summarizes the score of all three types of association (co-gained, co-lost, and gained-lost). The two vertical dotted lines at scores −5 and −6.8 correspond to FDRs of 0.05 and 0.005, respectively.

The false discovery rate (FDR) is the fraction of pairs at a given association score *t* for which this score is likely due to chance alone. It can be calculated as the number of pairs showing an equal or stronger association than *t* in the empirical data, divided by the corresponding number for the null model. Here, we examined associated gene pairs at FDR 0.05, corresponding to an association score of *t*=-5, and at FDR 0.005, corresponding to a score of *t*=-6.8 (see the two vertical dotted lines in Figure 1).

To test whether or not the associated gene pairs identified are indeed functionally connected, we examined the 8664 significantly associated pairs (at FDR 0.05; Supplemental Table S4) for which both genes have GO annotations (Gene Ontology Consortium 2015). We performed binomial tests to see if both share at least one identical GO term more often than expected by chance. Co-gained and co-lost pairs are significantly more likely to share the same GO term than random pairs (*p*<10^−6^ in each case, Table 1); however, this is not the case for gained-lost pairs (*p*=0.98, Table 1), suggesting that these pairs tend to be false positives.

**Table 1.** Statistical tests for functional relationships between associated gene pairs (FDR 0.05).

|  |  |  |  |
| --- | --- | --- | --- |
| | | | $N_{\text{success}} / N_{\text{trial}}^1, (p\text{-value})^2$ |

| Association type | total pairs | both have GO annotation | distance < 30kb | distance > 30kb | total |
| --- | --- | --- | --- | --- | --- |
| co-gained | 7538 | 2206 | 377 / 1581<br>(0.00016) | 248 / 625 (<1E-6) | 625 / 2206<br>(<1E-6) |
| gained-lost | 71 | 18 | - | 1 / 18<br>(0.9825) | 1 / 18<br>(0.98) |
| co-lost | 1055 | 612 | 164 / 419<br>(<1E-6) | 35 / 193 (0.7791) | 199 / 612<br>(<1E-6) |
| overall | 8664 | 2836 | 541 / 2000<br>(<1E-6) | 284 / 836 (<1E-6) | 825 / 2836<br>(<1E-6) |

<sup>1</sup> the null expectation is $N_{\text{success}} / N_{\text{trial}} = 20.12\%$
<sup>2</sup> $p$ -values from binomial test

### Distance distribution between co-gained gene pairs

We estimated the distance of co-gained gene pairs at the time that they were added to their ancestral host. In many cases, parts of horizontally transferred DNA segments will be lost subsequently, thereby reducing the distance between two co-transferred genes. Conversely, later genomic rearrangements may increase the original genomic distances. To estimate the genomic distance during the HGT event, we thus took the minimum distance between orthologs of the two co-gained genes in any of the examined genomes.

Figure 2 shows the cumulative distribution of the genomic distance for co-gained gene pairs selected at FDR 0.05 (solid lines) and 0.005 (dashed lines; see Supplemental Figure S4 for the corresponding probability density functions). Both curves show a pronounced kink at 30kb, indicating the existence of two stacked distributions. The dominant distribution at low genomic distances to the left of the kink likely represents the pairwise distances of genes that were co-gained in one HGT event; thus, 30kb appears to be an upper bound on the size of successfully transferred DNA segments. We also examined alternative definitions for the genomic distance of co-gained pairs, including the mean, median, and maximum of the distances between their two orthologs in any of the 53 extant genomes (Supplemental Figure S5). These alternative definitions, except for the one using maximal genomic distances, also show a corresponding kink at around 30kb. We conclude that the kink in Figure 2 is not an artifact of our definition of genomic distance for co-gained gene pairs, but that the length distribution of co-gained pairs is indeed bimodal.

**Figure 2.**
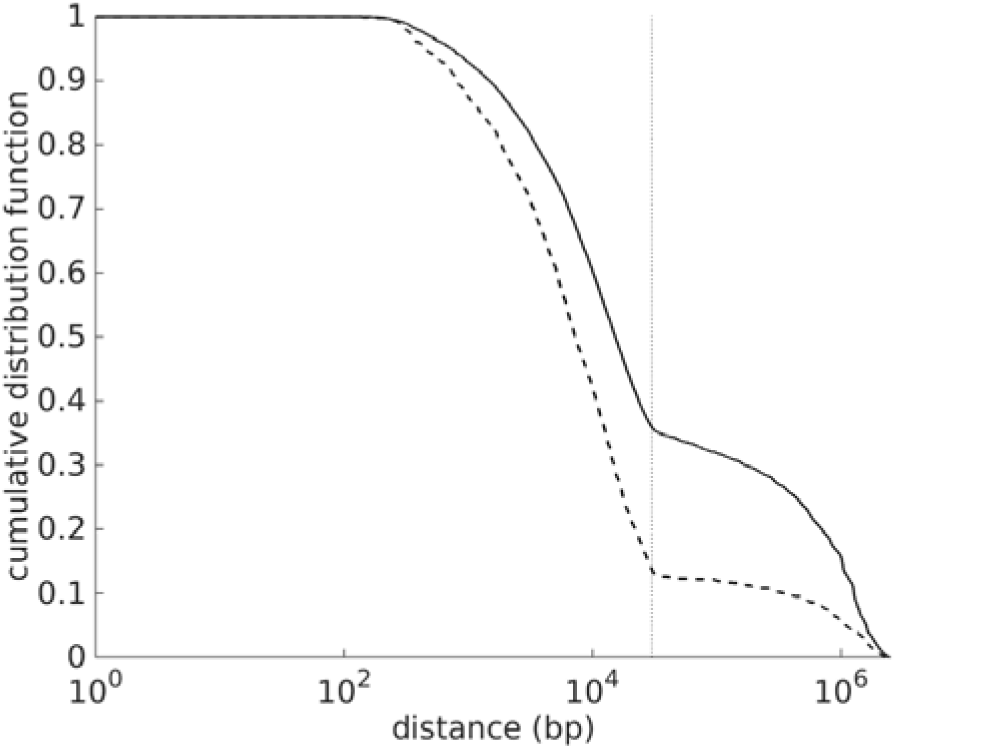
Cumulative distribution function of genomic distances between co-gained gene pairs at FDR 0.05 (solid line) and 0.005 (dashed line), showing a pronounced kink at a genomic distance of 30kb (vertical dotted line). See Supplemental Figure S4 for the corresponding probability density function, and Supplemental Figure S6 for the distribution of co-lost and gained-lost associations.

Pairs in the long-range mode of the distribution often display much larger distances. These pairs may in part correspond to the expected 5% of false positive co-gained pairs. However, these more distant pairs may also stem from independent gains of co-functioning genes in separate HGT events or from overestimates of the gene pairs’ ancestral genomic distances. Consistent with the latter two hypotheses, we found statistically significant support for functional relatedness of co-gained pairs not only at distances smaller than 30kb, but also at larger distances (Table 1).

The cumulative distance distribution for co-lost pairs is qualitatively very similar to that of the co-gained pairs, with a short-range component that ends at a kink at around 30kb (Supplemental Figure S6, green curve). However, the long-range component of the pairwise distance distribution of co-lost pairs is shifted downwards relative to the co-gained pairs (Supplemental Figure S6). This might be due to fewer overestimates of gene distances. HOwever, it would also be consistent with fewer independent gene losses of co-functioning genes; indeed, we found no enrichment of functionally related co-lost pairs at distances larger than 30kb. Thus, at least at the temporal resolution of the phylogeny in Supplemental Figure S1, co-losses of genes may be a largely mechanistic rather than a predominantly functional phenomenon.

In contrast to the pairwise distance distribution of co-gained and co-lost genes, the distribution for gained-lost associations (non-homologous replacements) appears to consist of only a single long-range component, showing no sign of a kink (Supplemental Figure S6, red curve). This observation is consistent with the rareness of significant pairwise gene associations: at FDR 0.05, there are 7,538 co-gained pairs and 1,055 co-lost pairs, but only 71 gained-lost pairs. In addition, the gained-lost pairs are not functionally related (Table 1). These findings suggest that the identified gained-lost associations predominantly represent noise and are not genuine non-homologous replacements.

### The distance distribution of co-gained genes cannot be explained by operon sizes

The short-range mode of the pairwise distance distribution of co-gained genes likely represents pairs that were acquired together in a single HGT event. Two evolutionary processes can shape this distance distribution. First, DNA from another organism must be taken up by the recipient. Thus, the maximal genomic distance of two co-transferred genes is limited by the carrying capacity of the transfer agent. Second, the vast majority of the analyzed gene gains are ancient. As bacteria tend to lose DNA that does not contribute to fitness (Batut et al. 2014), any co-gained gene pairs that persisted for such times are likely co-functional. Thus, provided that the carrying capacity of the transfer agent is sufficiently large, it is possible that the distance distribution of co-gained genes is dominated by the genomic organization of functional relationships.

It has even been suggested that natural selection favours the organization of functionally related genes into “selfish” operons to facility HGT (Lawrence and Roth 1996). If operons are indeed the basic unit of HGT, then the pairwise distance distribution of co-gained gene pairs ought to reflect the distribution of pairwise gene distances in operons. To test this prediction, we compared the short-range mode of the observed distance distribution with a pairwise distance distribution of genes generated from the known operons of *E. coli* K-12 MG1655 (Mao et al. 2009). Specifically, we modelled a superposition of a short-range distribution and a long-range distribution, where we set the short-range distribution to be the pairwise distribution of distances in operons, and the long-range distribution to be that of randomly chosen genes in the *E. coli* K-12 MG1655 genome. The model has one free parameter, 0 ≤ *w* ≤ 1, which defines the relative weight between the short and long-range distributions, but does not affect the shape of the two modes or the position of the boundary between them (Methods). We found that the model distribution (Figure 3, best fit at *w* = 0.65, dashed line) is unable to fit the empirical distribution (Figure 3, solid line). We conclude that the distances between gene pairs within *E. coli* operons are too short to explain the distance distribution of the observed pairs of co-gained genes.

**Figure 3.**
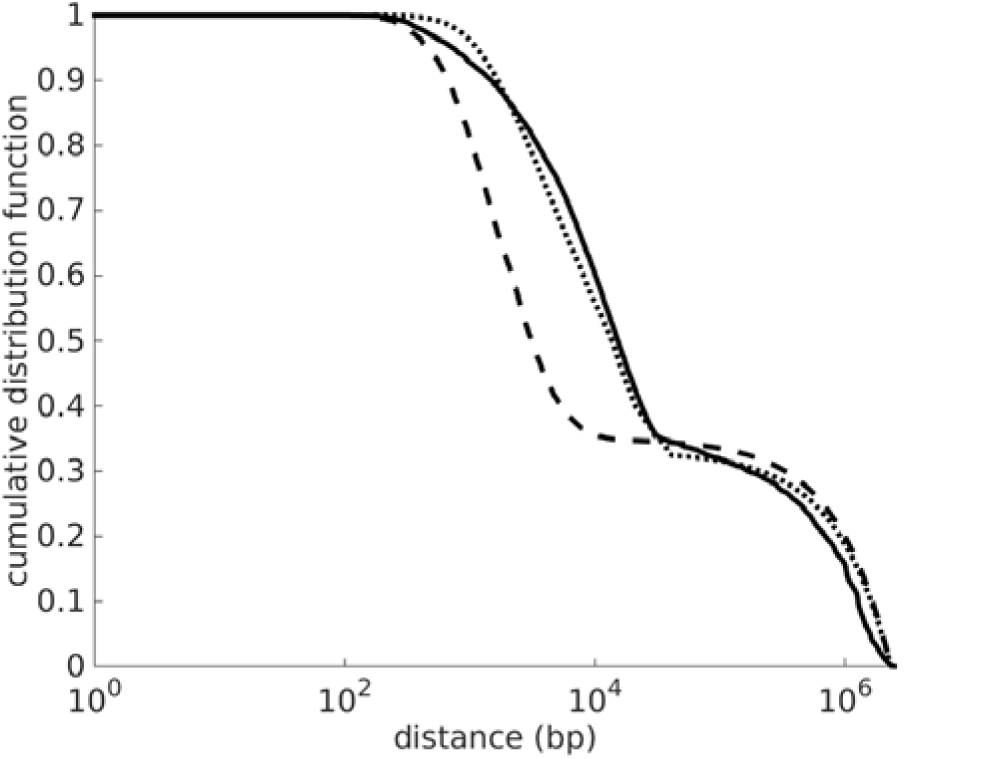
Comparison of the observed cumulative distance distribution of co-gained gene pairs (solid line) with the expectations from the operon model (dashed line) and the uber-operon model (dotted line). The operon model uses the distances of gene pairs in *E. coli* K-12 MG1655 operons (Mao et al. 2009) to represent the short-range distribution *ƒ*_*s*_(*x*) (best fit: *w* = 0.65). The uber-operon model uses the normalized functional autocovariance of *E. coli* K-12 MG1655 genes (best fit: *w* = 0.67) for the short-range distribution. See Supplemental Figure S7 for the corresponding probability density functions.

### Uber-operons can explain the distances of co-gained genes

Thus, to explain the distances between transferred gene pairs, we have to look at structures of functionally coupled genes that extend beyond operons. Such larger functional clustering units have been reported anecdotally and have been named uber-operons (Lathe et al. 2000); however, such structures are not well defined and were not investigated systematically.

We approximated the gene distance distribution in uber-operons through the functional autocovariance within a genome, defined as the probability for two nucleotides at a given genomic distance to be located in two different genes that have overlapping GO annotations (Gene Ontology Consortium 2015). We rescaled the autocovariance such that curves derived from different annotations (e.g., GO (Gene Ontology Consortium 2015) and InterPro (Hunter et al. 2009)) become comparable, and then normalized this rescaled autocovariance to obtain an approximation of the pairwise distance distribution of genes in uber-operon (Methods).

The rescaled functional autocovariance of all genes in *E. coli* K-12 MG1655 (Figure 4, solid line) drops to zero at around 25kb. This distribution is very similar to the autocovariance calculated only for regions harboring co-gained gene pairs (Figure 4, dashed line), indicating that co-gained gene pairs do not belong to unusual uber-operon structures.

**Figure 4.**
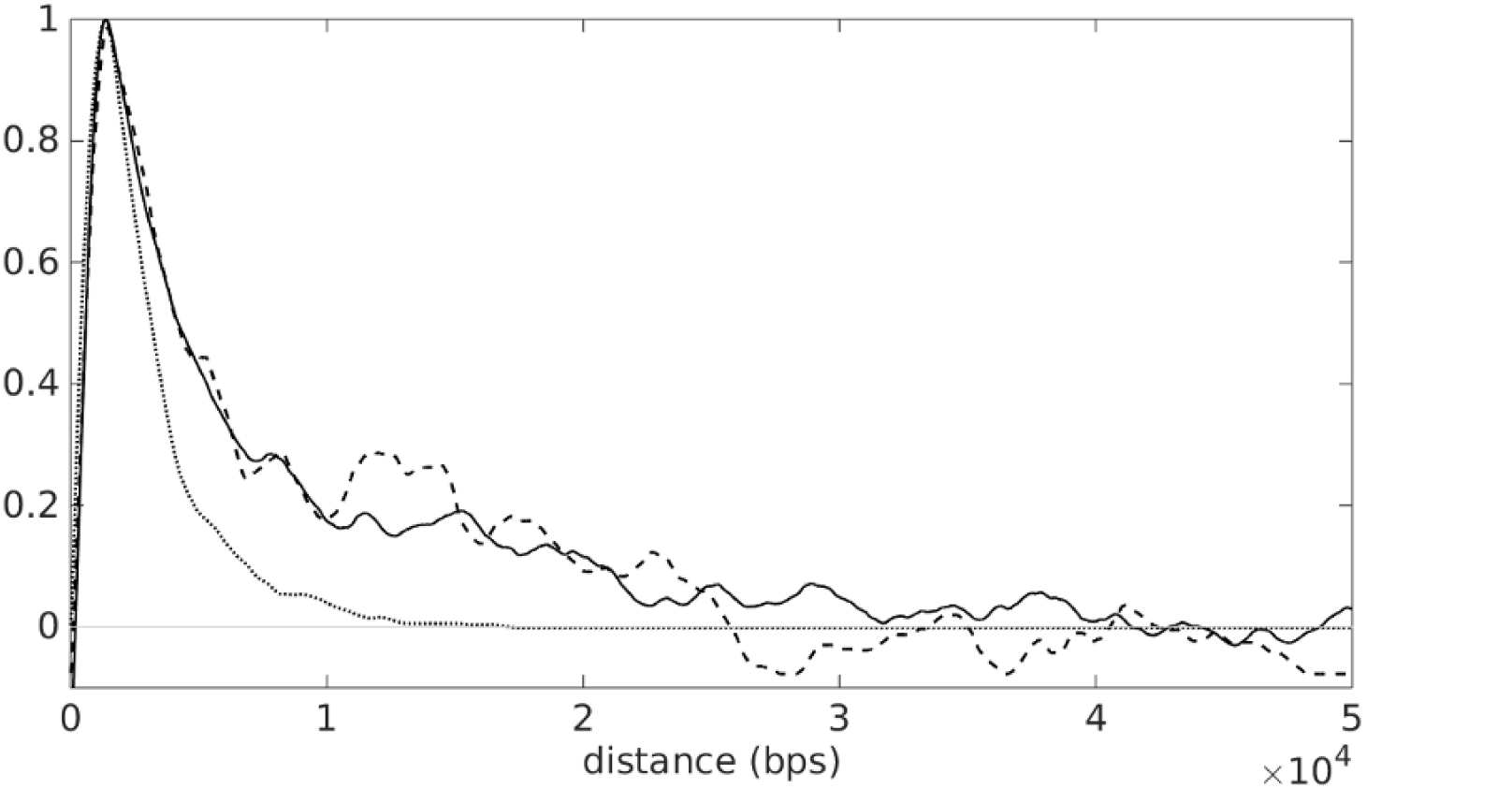
Rescaled functional autocovariance of genes in *E. coli* K-12 MG1655. The solid curve is calculated from all genes in *E. coli* K-12 MG1655, while the dashed curve is calculated only from genes that were consistently co-gained across the *E. coli* phylogeny according to our study. For comparison, the dotted line shows the functional autocovariance calculated only within *E. coli* K-12 MG1655 operons (Mao et al. 2009). At small distances, all three curves rise from small values to 1, as they leave the first gene (where autocovariance is 0 by our definition) and enter the second gene. These rescaled functional autocovariance curves then slowly decay to zero, indicating the finite size of uber-operons or operons. The operon model curve (dotted) decays substantially faster than the empirical functional autocovariance curves (solid and dashed), indicating that real functional clusters in *E. coli* K-12 MG1655 extend far beyond its operons.

To validate our estimation strategy for the distance distribution of functionally related genes, we also calculated the rescaled functional autocovariance restricted to positions within the same *E. coli* K-12 MG1655 operon (Mao et al. 2009) (dotted line); as expected, the normalized autocovariance curve results in a distribution that is very similar to the pairwise distance distribution of genes within operons (Supplemental Figure S3), confirming the validity of our approach. The operon-specific normalized autocovariance decays significantly faster than the overall autocovariance, with a cut-off at around 10kb that reflects the limited size of operons. This observed difference between the autocovariance within and beyond operons confirms that functional genomic clusters in *E coli* indeed often extend well beyond individual operons.

To see if the genomic distance distribution of functionally related genes beyond operons can explain the observed distance distribution of co-gained genes, we again constructed a model as a superposition of a short-range distribution (reflecting physical co-transfers) and a long-range distribution (reflecting independent transfers and false positives). This time, we used the estimated genomic distance distribution within uber-operons (the normalized autocovariance) to approximate the short-range mode of the empirical distribution. As shown in Figure 3 (best fit at *w* = 0.67, dotted curve), the predicted and empirical distance distributions of co-gained gene pairs are indeed highly consistent when basing the prediction on uber-operons instead of operons.

## DISCUSSION

We applied a simple statistical method to identify gene pairs repeatedly co-gained among 53 strains of *E. coli;* as expected, a comparison of the GO annotations (Gene Ontology Consortium 2015) shows that consistently co-gained gene pairs tend to be functionally related. Assuming that the majority of these gene pairs were transferred in the same HGT event, we estimated the distances of such pairs at the time they were co-transferred. Their distance distribution is bimodal, with a short-range mode that extends up to 30kb and a long-range mode that corresponds to the distance distribution of random gene pairs. The short-range mode likely represents true physical co-transfers.

There are three competing explanations for gene pairs in the long-range mode: (i) the genes in those pairs were transferred in independent HGT events; or (ii) they were transferred together but separated through subsequent genome rearrangements, such that we overestimated their ancestral distance. A detailed analysis of genome organization across E. coli strains could potentially distinguish these scenarios, but we leave this to future work.

The sharp 30kb upper bound for the short-range distribution of transferred gene pairs is intriguing: what constrains the size of transferred segments? An upper limit on the size of transferred DNA segments may be caused by the limited carrying capacity of transfer agents. The main mechanisms of HGT in *E. coli* are transduction and conjugation (Ippen-Ihler and Minkley 1986; Golomidova et al. 2007). Moreover, most phage genomes in the EMBL database have a sequence length well above 30kb (Kanz et al. 2005). Thus, the observed 30kb limit on successful co-transfers is too short to be explained by a limited carrying capacity of *E. coli*’s main HGT agents.

HGT consists of two stages: a mutational event that integrates a DNA segment into the host genome, followed by natural selection on the retention or removal of the segment or parts of it. Above, we argued that the short-range mode of the distance distribution cannot be explained by limitations on the mutational stage of HGT. Thus, we expected that the observed distribution instead reflects the distance distribution of co-functioning genes within the transferred DNA segments: only co-functioning genes are likely to be retained together by natural selection. To explain this short-range mode of the distance distribution, we initially hypothesized that it is consistent with the distances of gene pairs in operons, as these are often considered to be the basic unit of gene transfer (Lawrence and Roth 1996). Our simulations show that the distance distribution of co-transferred gene pairs expected from operon sizes underestimates the spatial extent of the observed short-range mode substantially (Figure 3). Instead, we found that the distance distribution of gene pairs in uber-operons (functional gene clusters beyond operons) is highly consistent with the observed distribution of co-transferred gene pairs.

Thus, we propose that uber-operons, rather than operons, are the basic units of HGT. In this work, we defined uber-operons based on functional autocovariance; previous work has linked such structures to co-regulated operons, regional variation in codon bias, and chromosomal coiling (Lathe et al. 2000; Warren and ten Wolde 2004; Hershberg et al. 2005; Touchon and Rocha 2016; Fritsche et al. 2011; Bailly-Bechet et al. 2006).

Interestingly, a recent analysis of domesticated phages revealed that while recently added phage segments can span larger distances, anciently domesticated phage segments also follow a distribution bounded by 30kb, likely because evolution has trimmed down such segments to genes useful to the host (Bobay et al. 2014). An analysis of the size of co-transferred DNA segments in *E. coli* based on regional codon bias also found an upper size limit of approximately 30kb (Bailly-Bechet et al. 2006). However, the same study also found a much larger upper limit for the size of regional codon bias in *Bacillus subtilis*, extending up to approximately 180kb (Bailly-Bechet et al. 2006). One possible explanation for this disparity is that uber-operons in *B. subtilis* are much larger than in *E. coli*. Alternatively, in contrast to *E. coli*, successful HGT in *B. subtilis* may not be dominated by the uptake of beneficial gene sets. While we restricted our analysis to the acquisition and loss of genes, the analysis of regional codon bias as performed for *B. subtilis* also includes the exchange of functionally identical sequences through recombination. In such cases, natural selection on gene retention plays no important role in determining the fate of horizontally transferred sequences; thus, in contrast to *E. coli*, the observed size distribution of transferred DNA segments in *B. subtilis* may be determined not by natural selection but by limits on DNA uptake via transformation.

In sum, we have shown that genes consistently co-transferred across *E. coli* strains follow a distance distribution that is consistent with uber-operons, i.e., functional gene clusters that extend beyond operons, as the basic unit of HGT. While higher-level functional clustering in bacterial genomes has been reported previously (Lathe et al. 2000; Warren and ten Wolde 2004; Hershberg et al. 2005; Touchon and Rocha 2016; Fritsche et al. 2011; Bailly-Bechet et al. 2006), these structures have so far not been linked to HGT. Future studies on the properties and the evolution of uber-operons may greatly contribute to our understanding of the structure and evolution of prokaryotic genomes.

## METHODS

### Reconstruction of the phylogenetic tree

To infer HGT, we first needed to establish a species tree reflecting vertical inheritance. We obtained the genbank files for 53 *E. coli* strains and 17 sequences of closely related species (Supplemental Table S1) from NCBI (NCBI Resource Coordinators 2013; Benson et al. 2009). We extracted the amino acid sequences of all genes and identified orthologous gene groups using Proteinortho (Lechner et al. 2011) with the synteny option. We identified a total of 16,264 orthologous gene families in the 53+17 strains (Supplemental Table S2 lists the proteinortho results that maps the orthologous gene families to the genes in the strains, and Supplemental Table S3 lists the gene names, locus tags, gene IDs and protein IDs for each orthologous gene family).

The amino acid sequences of the 1,334 one-to-one orthologs universal to all 70 genomes were aligned using MAFFT (Katoh and Standley 2013) with default parameters. We then concatenated the alignments and estimated a phylogeny of vertical inheritance for these 70 genomes using RAxML (Stamatakis 2014) with 200 fast bootstraps and with the “PROTCATAUTO” option for model choice. This protocol generated a phylogenetic tree with at least 60% bootstrap support at each internal branch.

The phylogenetic tree was rooted to group all 53 *E. coli* strains into a monophyletic subtree, with each of the 52 internal nodes of this subtree considered as an ancestral strain (see Supplemental Figure S1 for the phylogenetic tree of the 53 *E. coli* strains, Supplemental Figure S2 for the tree of all 70 strains, and Supplemental Data File D1 for the Newick format of the 70 strain tree).

### Reconstructing ancestral genomes and inferring the genes acquired through HGT

We used the GeneTRACE maximum parsimony algorithm (Kunin and Ouzounis 2003) to determine the presence and absence of each gene at the 52 ancestral nodes, based on its presence and absence on the 53 extant genomes. Note that we preferred not to use a maximum-likelihood method to infer ancestral genome content, as existing methods assume constant rates of gain and loss along the phylogeny for each gene; this is unlikely to reflect evolutionary history, and often leads to the inference of multiple gains and losses on a single branch.

If a gene is present in the ancestral node, but absent in the descendant node, then we designate it as lost on the corresponding branch of the phylogenetic tree. If a gene is absent in the ancestral node of a branch but present in the descendant node, then we designate it as gained on the corresponding branch (see Supplemental Table S5 for the orthologs in each of the extant and ancestral strains).

### Identifying the evolutionary associations between gene pairs

For each of the 16,264 orthologous gene families, we represented the gain and loss history along the 104 branches of the phylogenetic tree by two separate binary vectors of 104 elements: if a gene is gained in an evolutionary step (i.e., on one branch), then the corresponding element in its gain-vector is 1, and 0 otherwise; if it is lost in a step, then the corresponding element in its loss-vector is 1, and 0 otherwise.

Next we quantified pairwise evolutionary associations between genes. For each pair of vectors, we summed the occurrence of the four element-wise patterns (0,0), (0,1), (1,0) and (1,1) over the 104 rows and represented the sums as a 2-by-2 contingency table. Further, as each gene has two vectors representing its gain and loss history, there can be three types of associations between two gene families A and B: (i) co-gain of gene A and gene B on the same branch; (ii) gain of A with loss of B, or loss of A with gain of B; and (iii) co-loss of A and B. The association score of a pair of vectors is defined as the decadic logarithm of the *p*-value at the right tail of Fisher’s exact test.

### Null model of gene association

We defined the score of association between gene pairs in the empirical data based on *p*-values from Fisher’s exact test, but these values have no straight-forward statistical interpretation. This is because Fisher’s exact test assumes that observations are independent of each other, but the gain and loss patterns of genes on different branches are not: e.g., when a gene is gained in one step, it cannot be gained again in the subsequent step. We also developed a null model of gene transfer, where the presences and absences of each gene among different extant strains is randomly shuffled. The same algorithm of maximal parsimony was then applied to reconstruct randomized ancestral genomes, and the association scores between genes in this null model were calculated.

We defined the false discovery rate (FDR) to describe the significance of association by comparing the distribution of association score between the empirical data and the null model. Let *N*_*d*_(*t*) and *N*_*n*_(*t*) be the number of gene pairs with association scores more significant than *t* in the empirical data and null model, respectively; a gene pair with score *t* then has 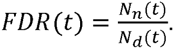. Figure 1 shows that an FDR of 0.05 (0.005) corresponds to a score of −5.0 (-6.8).

### Assigning GO categories through UNIPROT

We queried the UNIPROT database (U**niProt Consortium 201**5) to obtain the protein entries that match our orthologous gene families. For each orthologous gene family, we extracted the gene name and locus tag of each of its corresponding genes from the Genbank files, and used them as keywords to query the database for entries that match these names and tags (Supplemental Table S3). We then filtered out the entries with organism names that do not contain “Escherichia coli” or “Shigella”; gene names or locus tags that return multiple entries can cause confusion and so their results were also ignored.

In the end, each orthologous gene family could map to more than one UNIPROT entry. The annotation of each orthologous gene family with gene ontology (GO) terms (Gene Ontology Consortium 2015) is then defined as the union of the individual entries.

### Modelling the distribution of gene pair distances

We developed a simple model to explain the distance distribution of the associated gene pairs. We assumed that a pair of associated genes can have one of two origins: (i) both gained in the same transfer event, and so their distance is described by a short-range distribution *ƒ*_*s*_(*x*), which reflects the limit of the transfer carrier or the distance distributions in source genomes; or (ii) both genes are transferred independently, and so their distance follows a long-range distribution *ƒ*_*l*_(*x*), which reflects the limit imposed by the host genome. We used a parameter 0 ≤ *w* ≤ 1 to specify the relative weight of the two modes (*w* is the fraction of gene pairs acquired in the same transfer event among all gene pairs designated as associated). Note that *w* does not affect the boundary between the two modes of the distribution or the shapes of the two modes.

We denoted the short-range distribution as *ƒ*_*s*_(*x*) and the long-range one as *ƒ*_*l*_(*x*). We fixed *ƒ*_*l*_(*x*) to be the pairwise distance distribution between all genes in *E. coli* K-12 MG1655; for *ƒ*_*s*_(*x*), we tested different pairwise distance distributions based on operons or on uber-operons (see main text). The overall distribution is

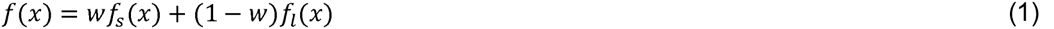

We determined *w* by fitting the empirical distribution of pairwise gene distances of associated pairs to Equation (1). Fitting was done by minimizing the area between the cumulative distributions of the empirical data and that of the model, assuming a logarithmic scale on the x-axis.

### Delineation of uber-operons through functional autocovariance

The functional autocovariance *G*(*x*) can be used to measure the extent of clusters of functionally coupled genes. Specifically, given a nucleotide at site *i* = 0 within a gene with a given set of GO annotations (Gene Ontology Consortium 2015), *G*(*x*) measures the probability for another nucleic site *j* at distance *x* = *j* - *i* to be within a gene that has at least one GO annotation in common. Let us define *g*_*t*_(*x*) to be a discrete function that maps distance *x* to ones and zeros: *i* can be any nucleic site on the *E. coli* K-12 MG1655 genome within a gene that has at least one GO annotation; *x* is a positive integer; *g*_*t*_(*x*) = 1 if *x* and *x* + *i* are sites of two different genes sharing at least one GO term, and 0 otherwise. The functional autocovariance is then

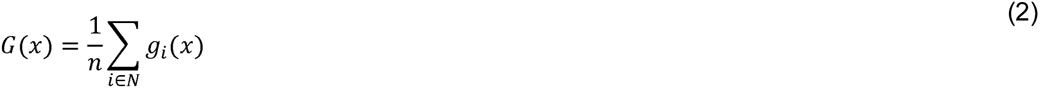

where *N* is the set of all nucleic sites of genes considered, and *n* is the number of elements in *N*.

We assume that there is a higher chance for gene pairs within the same functional cluster to share a GO annotation, but a lower chance for pairs across different functional clusters; thus, we expect *G*(*x*) to have the form

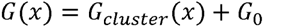

where *G*_*cluster*_(*x*) is the part of *G*(*x*) that is due to functional clustering, while *G*_0_ is the background probability that two random genes share a GO annotation. Gene pairs with small separation (small *x*) are likely to be in the same functional cluster and share the same annotation; but as *x* increases, this chance decays to *G*_0_.

Furthermore, while we here used the GO annotation to estimate the functional autocovariance, the same formalism can also be based on annotations other than GO, such as InterPro (Hunter et al. 2009). To make the functional autocovariance comparable across different sets of annotations, we rescale *G*(*x*) into 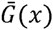, as

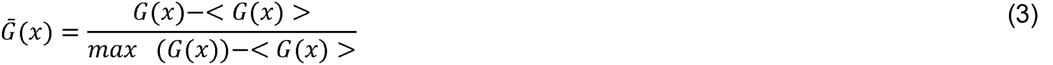

here, < *G*(*x*) >is the mean of *G*(*x*), while *max* (*G*(*x*)) is the maximum of *G*(*x*).

Regardless of the annotation used for the calculation of the autocovariance, the maximum of the rescaled autocovariance 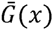 is 1, and decays to 0 at large distances. Once 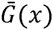 has decayed to zero, it will only represent noise. Our goal is to estimate the pairwise distance distribution of genes in uber-operons from 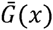; thus, we cut 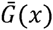 off at the point where it first crosses the x-axis beyond the bulk of the distribution (at *x* > 41410). To avoid any influence from unusually large genes, we did not score functional relationships of nucleotides within the same gene. This leads to an additional noise term at low distances, which we also removed (at *x* < 184) before using the resultant distribution to fit the distance distribution of co-gained genes. Normalization is then performed to ensure that the terms for all *x* sum to 1, which converts 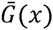 into an estimation of pairwise distance distribution of genes in function clusters.

To test the reliability of our approximation, we applied it also to gene pairs within operons in *E. coli* K-12 MG1655. Supplemental Figure S3 compares the distance distribution of gene pairs in K-12 MG1655 operons, and the distance distribution approximated using the rescaled and normalized autocovariance function of gene pairs in K-12 MG1655 operons that share GO annotations (with *x* > 17143 trimmed to remove noise). The plot shows that while the bulk of the distribution is slightly biased to the right, the right tail is well conserved.

## ACKNOWLEDGES

This work was supported by the German Research Foundation (DFG-grant CRC 680 to MJL).

